# *In vitro* reconstitution of chromatin domains

**DOI:** 10.1101/2023.02.27.530214

**Authors:** Kimberly Quililan, Elisa Oberbeckmann, Patrick Cramer, A. Marieke Oudelaar

## Abstract

The spatial organization of the genome modulates nuclear processes, including transcription, replication, and DNA repair^1,2^. Eukaryotic genomes are organized into distinct 3D chromatin domains^3^. However, the molecular mechanisms that drive the formation of these domains are difficult to dissect *in vivo* and remain poorly understood. Here, we reconstitute *S. cerevisiae* chromatin *in vitro* and determine its 3D organization at sub-nucleosome resolution by MNase-based chromosome conformation capture and molecular dynamics simulations. We show that regularly spaced and phased nucleosome arrays form chromatin domains *in vitro* that resemble domains *in vivo*. This demonstrates that neither loop extrusion nor transcription are required for domain formation. In addition, we find that the boundaries of reconstituted domains correspond to nucleosome-free regions and that insulation strength scales with their width. Finally, we show that domain compaction depends on nucleosome linker length, with longer linkers forming more compact structures. Together, our results demonstrate that nucleosome positioning is sufficient to reconstitute chromatin domains and provide a proof-of-principle for bottom-up 3D genome studies.

## INTRODUCTION

Eukaryotic genomes are organized into chromatin structures across different scales. The smallest unit of chromatin is the nucleosome core particle, which is formed by 147 base pairs (bp) of DNA wrapped around a histone octamer^4–6^. Nucleosome core particles are connected by short DNA “linkers” and form nucleosome arrays, which further organize into secondary structures that define the orientation of subsequent nucleosomes with respect to each other. Traditionally, nucleosome arrays were thought to form regular 30 nm fibers with a solenoid or zig-zag geometry^7^. However, more recent data based on X-ray scattering^8^, super-resolution light microscopy^9^, electron microscopy^10,11^ and innovative sequencing-based approaches^12^ have shown that *in situ* chromatin predominantly forms heterogenous and flexible secondary structures. At a larger scale, eukaryotic genomes organize into self-interacting chromatin domains. In mammals, chromatin domains are formed by at least two distinct mechanisms^2^. First, active and inactive regions of chromatin form functionally distinct compartments that span a wide range of sizes^13^. Second, a process of loop extrusion, mediated by Cohesin and CTCF, organizes the genome into local structures termed Topologically Associating Domains (TADs) that usually range from 100 kbp to 1 Mbp in size^14,15^.

The higher-order organization of the genome into chromatin domains is conserved in eukaryotes with smaller genomes, including *D. melanogaster*^16^ and *S. cerevisiae*^17,18^, in which domain sizes range from 10-500 kbp and 2-10 kbp, respectively. The nature of the chromatin domains in these species and the mechanisms by which they are formed are less well understood. Because the boundaries of chromatin domains in fly^16,19,20^ and yeast^18,21^ frequently overlap with promoters of highly transcribed genes, it has been proposed that the process of transcription or the transcriptional state of chromatin are key determinants of chromatin organization. There is currently no conclusive evidence for cohesin-mediated loop extrusion during G1 interphase in these species, but it is possible that this process also contributes to the basic organization of their genomes. Progress in our understanding of the conserved, core mechanisms that drive higher-order genome organization in eukaryotes is complicated by the close relationship and functional interplay between chromatin domains, chromatin state, and transcription *in vivo*. The fundamental principles underlying the formation of 3D chromatin structures are therefore difficult to disentangle and remain poorly understood.

The relatively recent development of inducible protein degradation approaches has been instrumental in identifying proteins and processes that are involved in the formation of higher-order chromatin structures^22^. However, even with these rapid protein depletion approaches it is challenging to determine the basic principles underlying the 3D organization of the genome, because the regulatory proteins involved often have overlapping or redundant functions that are difficult to dissect in the complex *in vivo* nuclear milieu. In addition, it is often not possible to distinguish direct from indirect effects following the perturbation of critical regulatory proteins, which limits conclusions about cause-consequence relationships.

To identify the molecular mechanisms that drive the formation of higher-order 3D chromatin structures in a controllable experimental set-up, we set out to reconstitute chromatin domains, using an *in vitro* system for yeast chromatin reconstitution. To map the spatial organization of individual reconstituted nucleosomes, we established a chromosome conformation capture (3C) approach to map folding patterns of *in vitro* chromatin at sub-nucleosome resolution. Using this unique approach, we show that reconstitution of regularly spaced and phased nucleosome arrays by the addition of purified transcription factors and ATP-dependent chromatin remodelers drives higher-order nucleosome folding into chromatin domains with remarkable similarity to *S. cerevisiae* genome organization *in vivo*. The boundaries of these domains correspond to the nucleosome-free regions (NFRs) at the transcription factor binding sites. We find that the strength of these boundaries depends on a combination of nucleosome array regularity and NFR width. Our work therefore shows that the formation of regular and phased nucleosome arrays is sufficient to reconstitute native chromatin domains in yeast and demonstrates that neither loop extrusion nor transcription are a prerequisite for domain organization. By comparing three different remodelers that set distinct nucleosome linker lengths, we also show that compaction of chromatin domains is dependent on the linker length. 3D chromatin models generated with molecular dynamics simulations confirm these results and highlight the fundamental principles underlying chromatin domain formation. Together, our work identifies an underappreciated function of nucleosome positioning in the formation of chromatin domains, which has important implications for our understanding of genome organization across eukaryotic species.

## RESULTS

### An *in vitro* system to study chromatin domain formation

We established an approach to study higher-order folding of *in vitro* reconstituted chromatin (Fig. 1a). To this end, we adapted a previously established system to reconstitute *S. cerevisiae* chromatin *in vitro*^23,24^. As a DNA template, we used a genomic plasmid library covering *S. cerevisiae* chromosomes V-IX. Each of these plasmids contains a ~7 kb backbone and an insert covering a fraction of the *S. cerevisiae* genome with an average length of ~10 kb. Incubating the plasmid library with purified recombinant *S. cerevisiae* histone octamers (Extended Data Fig. 1a) under high-salt conditions and dialyzing overnight into low-salt buffer leads to spontaneous assembly of nucleosomes. To obtain high nucleosome densities that resemble *in vivo* chromatin^25^, we used negatively-supercoiled plasmid DNA amplified in *E. coli*, as it is thought that negative supercoiling propagates nucleosome assembly during salt gradient dialysis (SGD)^26,27^.

**Figure 1.**
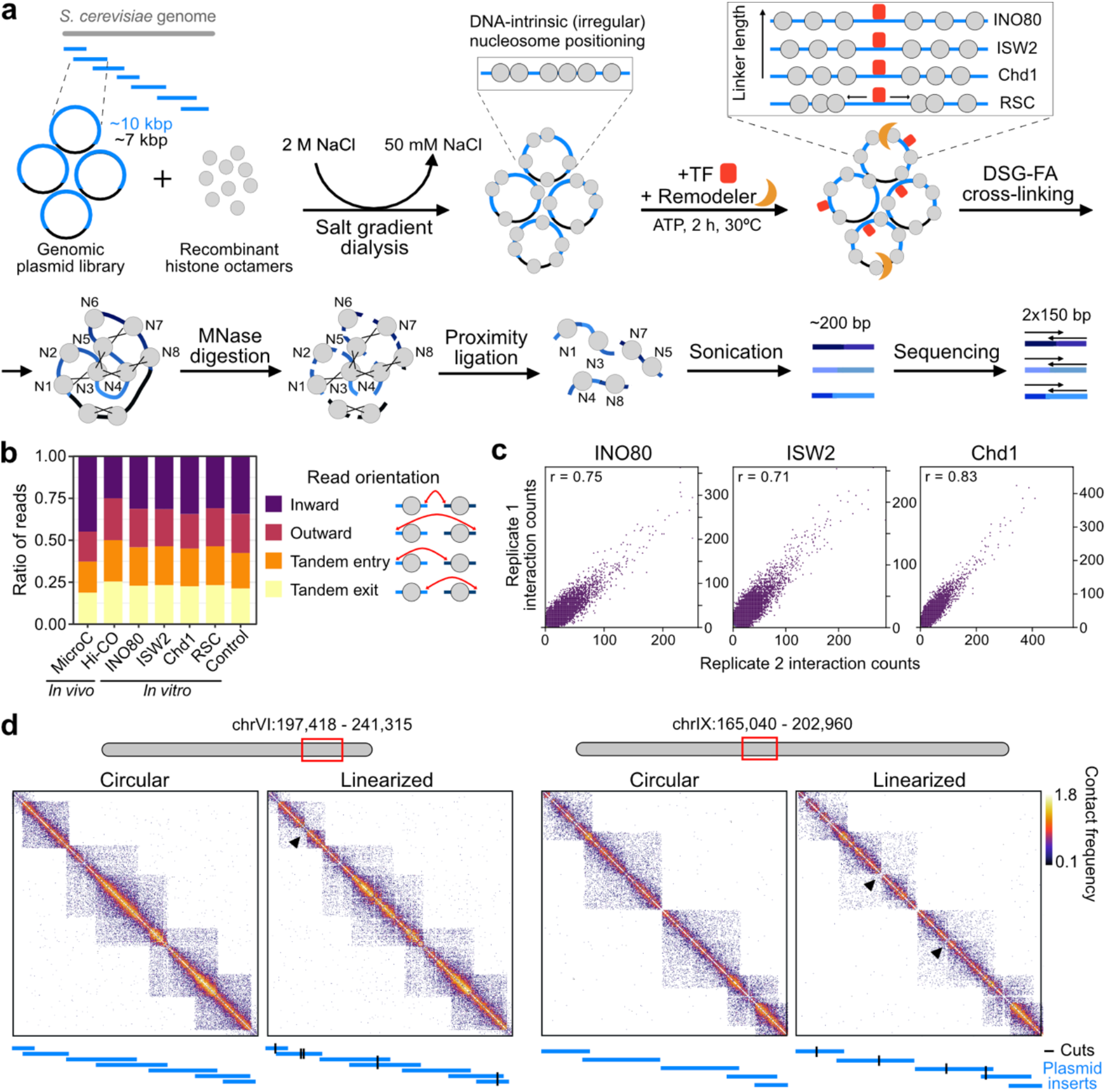
An in vitro system to study higher-order chromatin structure. **a,** Schematic overview of in vitro Micro-C. TF: transcription factor; DSG: disuccinimidyl glutarate; FA: formaldehyde. **b,** Proportion of read numbers clustered by the orientation of the interacting nucleosomes. For comparison, in vivo Micro-C^21^ and Hi-CO^39^ data are shown. **c,** Correlation plots of interaction frequencies of in vitro Micro-C replicates. Pearson’s correlation coefficient r is indicated. **d,** Contact matrices of two different genomic regions of in vitro Micro-C. SGD chromatin was incubated with transcription factors and INO80 (circular) or with transcription factors, INO80 and the restriction enzyme BamHI (linearized). Plasmids are shown in blue; BamHI cut sites are shown in black; arrowheads show sites with strong BamHI cleavage which leads to separation of the domains. Log^10^ interaction counts are plotted at 80 bp resolution.

The positioning of nucleosomes in SGD chromatin is solely directed by the DNA sequence and therefore irregular. To reconstitute regular nucleosome positioning, we incubated the SGD chromatin with purified, sequence-specific DNA-binding transcription factors and ATP-dependent chromatin remodelers (Extended Data Fig. 1a). As transcription factors, we used the General Regulatory Factors (GRFs) Abf1 and Reb1. These factors bind promoter regions and act as pioneer factors and transcriptional activators^28–30^ (Extended Data Fig. 1b). As chromatin remodelers, we used INO80, ISW2, and Chd1. These remodelers create regular nucleosome arrays at transcription factor binding sites that resemble *in vivo* chromatin (Extended Data Fig. 1b). The nucleosomes in these arrays are evenly spaced and phased relative to reference sites; in our system these correspond to Abf1 and Reb1 binding sites (Extended Data Fig. 1b). INO80, ISW2, and Chd1 have an intrinsic “ruler” function that determines the spacing between the nucleosomes in the reconstituted arrays. INO80 generates relatively large linkers, ISW2 forms medium-sized linkers, and Chd1 forms relatively small linkers^23,31^. We also used the chromatin remodeler RSC, which does not have spacing activity, but plays an important role in maintaining NFRs (Extended Data Fig. 1b)^30,32^. For our experiments, we incubated the SGD chromatin with both transcription factors and one of the remodelers. As a control we used chromatin incubated with the two transcription factors only.

To analyze the 3D folding patterns of reconstituted chromatin, we established a 3C approach that is compatible with our *in vitro* set-up. 3C is based on digestion and subsequent proximity ligation of crosslinked chromatin and thereby allows for detection of spatial proximity between DNA sequences using high-throughput sequencing^33–35^. An important aim of our study is to identify the folding pattern of individual nucleosomes, which requires 3C analysis at very high resolution. The recent development of the Micro-C^18,21,36–38^, Hi-CO^39^, and Micro-Capture-C (MCC)^40,41^ approaches has demonstrated that the use of micrococcal nuclease (MNase) instead of restriction enzymes for chromatin digestion significantly increases the resolution of 3C data. Since the MCC approaches allow for direct sequencing of ligation junctions in regions of interest, they support a resolution of 1-20 bp^40,41^. We therefore adapted the MCC protocol to enable high-resolution analysis of the 3D conformation of SGD chromatin (Fig. 1a). Instead of using capture oligonucleotides to enrich for a subset of the yeast genome, we used a plasmid library that covers only chromosomes V-IX, approximately a quarter of the yeast genome. This corresponds to ~3 Mb of DNA in total, which can be sequenced with high coverage and thus analyzed at very high resolution.

To enable efficient 3D analysis of SGD chromatin, we optimized the MCC crosslinking, digestion, and ligation conditions (Methods). We refer to the optimized procedure as *in vitro* Micro-C. In all experiments, we used a small fraction of the digested sample for MNase-seq^42^ to confirm efficient remodeling and regular *in vivo*-like nucleosome positioning (Extended Data Fig. 1b). In the conventional MCC protocol, the 3C procedure is performed in intact cells that have been permeabilized with digitonin and crosslinked with the short-range crosslinking agent formaldehyde (FA). However, we find that we are not able to efficiently detect long-range interactions between nucleosomes in chromatin that has been crosslinked with FA, irrespective of the FA concentration used (Extended Data Fig. 2a-c; Supplementary Table 1). It has previously been shown that the addition of the long-range crosslinking agent disuccinimidyl glutarate (DSG) improves the efficiency with which interactions that span long genomic distances are captured in the Micro-C procedure in spheroplasts^21^. We therefore tested “double crosslinking” conditions with different FA and DSG concentrations and find that this drastically improves our ability to identify long-range interaction patterns (Extended Data Fig. 2a-c; Supplementary Table 1). We speculate that long-range crosslinking in an *in vitro* set-up is critical, given the lack of a protein-dense stabilizing chromatin environment that is present in nuclei *in vivo*. Importantly, nucleosome positioning as measured by MNase-seq is not affected by the crosslinking method (Extended Data Fig. 2d). We therefore implemented double crosslinking in the *in vitro* Micro-C protocol and adapted the MNase concentration and ligation conditions to achieve efficient digestion and re-ligation (Methods, Extended Data Fig. 2e). After reverse crosslinking and purifying the ligated DNA, we sheared the DNA to ~200 bp and performed 150 bp paired-end sequencing. Since the *in vitro* Micro-C procedure allows for direct identification of the ligation junctions, it is possible to define the orientation of the interacting nucleosomes based on the direction of the reads and to distinguish reads resulting from an inward interaction and reads resulting from regions that have never been digested (Fig. 1 b; Methods). Similar to Hi-CO, the *in vitro* Micro-C protocol therefore allows for detection of nucleosome orientation without a significant bias towards detecting inwardly oriented nucleosomes (Fig. 1b).

Comparison of *in vitro* Micro-C data of independent samples of remodeled chromatin shows a high degree of correlation between replicates, thus confirming the robustness of the *in vitro* Micro-C protocol (Fig. 1c). Importantly, analysis of the interaction patterns of SGD chromatin shows a strong enrichment of intra-plasmid interactions compared to inter-plasmid interactions (Fig. 1d, Supplementary Table 1). This indicates that *in vitro* Micro-C predominantly captures meaningful higher-order interactions between nucleosomes contained within individual plasmids with minimal spurious ligation between different plasmids. To investigate whether these interaction patterns are influenced by topological constraints related to the circular nature of the plasmids, we also generated Micro-C data of linearized SGD chromatin. For these experiments, we added to the remodeling reaction the restriction enzyme BamHI, which has a 6 bp recognition motif and cuts on average on 1-2 sites per plasmid. As expected, we find that plasmid linearization creates additional interaction “boundaries” at the BamHI restriction sites, since the regions upstream and downstream are no longer adjacent but on opposite ends of the linear plasmid (Fig. 1d). However, importantly, linearization of the plasmid does not affect nucleosome positioning (Extended Data Fig. 2f) or patterns of higher-order nucleosome folding (Fig. 1d and Extended Data Fig. 2g,h).

### Regular nucleosome arrays form chromatin domains

To investigate the higher-order folding patterns of SGD chromatin in further detail, we plotted contact matrices of regions contained on individual plasmids at high resolution (40 bp). We excluded regions covered by multiple plasmids; the matrices therefore span approximately 3-7 kbp. Since chromatin domains in yeast are on average 2-10 kbp in size, these matrices allow for detailed comparison of domain organization in *in vitro* and *in vivo* chromatin^21^ (Fig. 2; Extended Data Fig. 3a).

**Figure 2.**
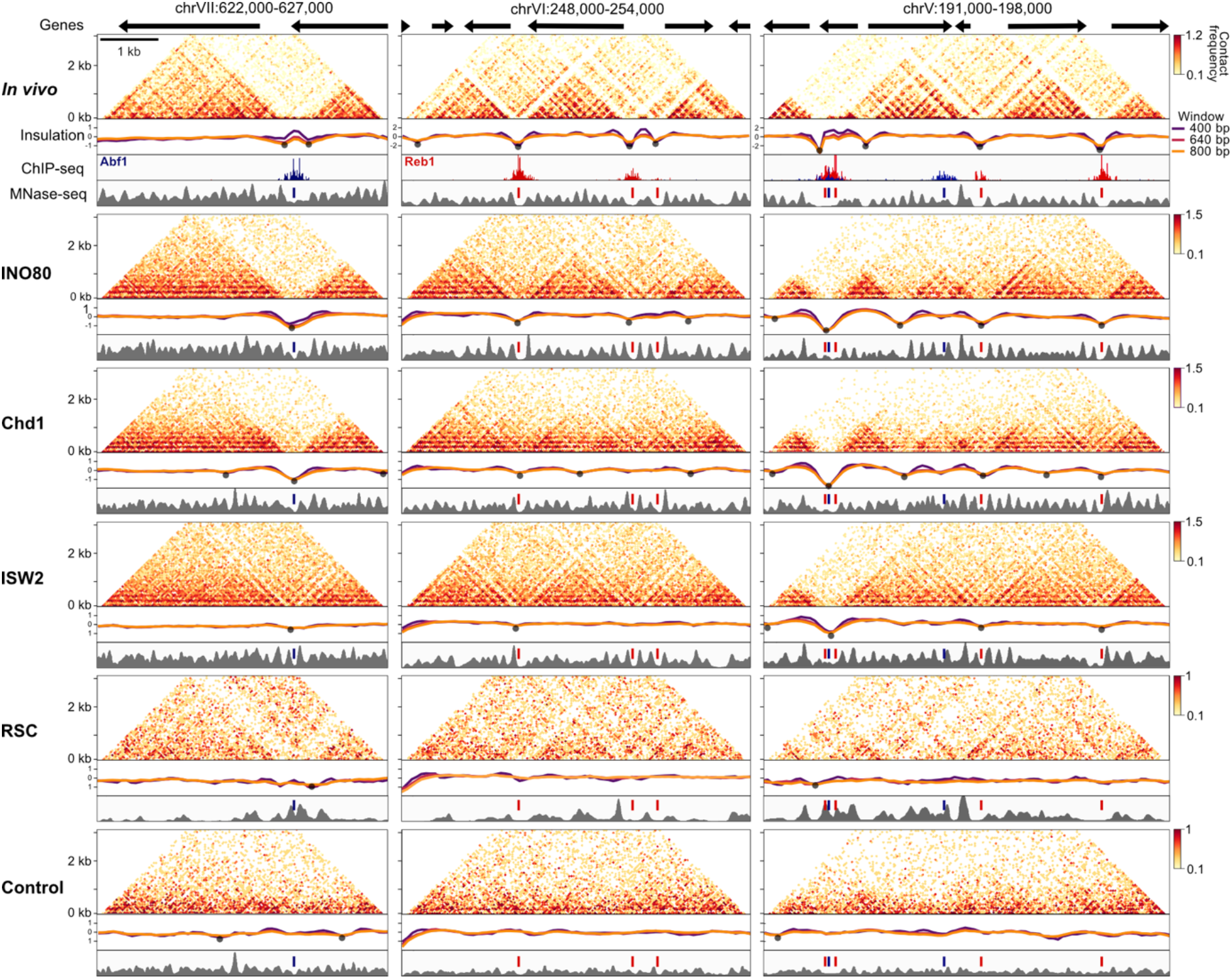
Regularly spaced and phased nucleosome arrays are sufficient to reconstitute chromatin domains. Contact matrices displaying in vivo^21^ and in vitro Micro-C data with corresponding insulation scores and nucleosome occupancy profiles (MNase-seq) plotted for three different genomic regions. Abf1 and Reb1 ChIP-exo^28^ data are shown below the in vivo data. Chromatin used for in vitro Micro-C was incubated with the indicated remodeler and the transcription factors Abf1 and Reb1 or with the transcription factors only (control). Strongly insulating boundaries are labeled with black dots. Blue and red lines in MNase-seq profiles indicate main Abf1 and Reb1 peaks, respectively. Micro-C data are plotted as log^10^ interaction counts at 40 bp resolution. Insulation scores are calculated at 80 bp resolution with three different sliding windows.

As we only use 2 out of 256 putative yeast transcription factors^43^, we focused our analyses on regions with a high abundance of Abf1 and Reb1 binding sites. These factors support the formation of regularly spaced and phased *in vivo*-like chromatin (Extended Data Fig. 1b). We find that the patterns of higher-order genome folding in such regions of *in vitro* chromatin are dependent on the chromatin remodeler used during reconstitution. Interestingly, the presence of chromatin remodelers that generate regular nucleosome arrays (INO80, Chd1, and ISW2) drives higher-order genome folding into chromatin domains with striking similarity to domain organization *in vivo*. The boundaries of these domains correspond to the NFRs that are formed at Abf1 and Reb1 binding sites, as confirmed by the corresponding nucleosome occupancy patterns derived from the MNase-seq data. However, reconstitution with a chromatin remodeler involved in NFR formation but without spacing activity (RSC) does not lead to the formation of *in vitro* chromatin domains. Similarly, reconstitution with transcription factors only in absence of a remodeler does not lead to specific 3D interaction patterns. We therefore conclude that nucleosome positioning directly influences higher-order genome folding and that the presence of regularly spaced and phased nucleosome arrays is required and sufficient for the formation of chromatin domains.

Although the occupancy and 3D organization of nucleosomes in *in vitro* chromatin in the presence of remodelers with spacing activity is remarkably similar to *in vivo* chromatin, there are subtle differences. Most notably, we observe the generation of additional NFRs in *in vitro* chromatin remodeled with INO80 and Chd1, which correspond to the formation of additional domain boundaries (Fig. 2; right panel). Interestingly, the underlying DNA sequences are enriched in features that are associated with the formation of particular DNA shapes, including helical twists and propeller twists^44^ (Extended Data Fig. 3b). It has previously been shown that these shapes can be recognized by some chromatin remodelers, including INO80^23,24^. We speculate that DNA shape has a more prominent role in regulating nucleosome occupancy and structure in the minimal *in vitro* system compared to the *in vivo* milieu, in which additional nucleosome positioning mechanisms might overrule the influence of DNA shape on chromatin organization.

### Insulation strength of domain boundaries depends on NFR width

To explore the relationship between Abf1 and Reb1 enrichment and the formation of domain boundaries in a genome-wide manner, we correlated the called boundaries (Fig. 2) with ChIP-exo data^28^ (Extended Data Fig. 4a). We find that regions with strong insulation are highly enriched for Abf1 and Reb1 binding sites, both *in vivo* and *in vitro*. To investigate this further, we performed pile-up analyses of the chromatin interactions in a 3 kbp region surrounding the transcription factor binding sites in the *in vivo* and *in vitro* Micro-C data (Fig. 3a, Extended Data Fig. 4b). The resulting *in vivo* “meta” matrices show three distinct interaction patterns: (1) a banding pattern parallel to the diagonal, which indicates regular spacing between nucleosomes; (2) horizontal and vertical bands that form a “grid” of phased nucleosomes that are aligned to the NFR at the Abf1 and Reb1 binding sites; (3) insulation between the regions upstream and downstream of the transcription factor binding sites.

**Figure 3.**
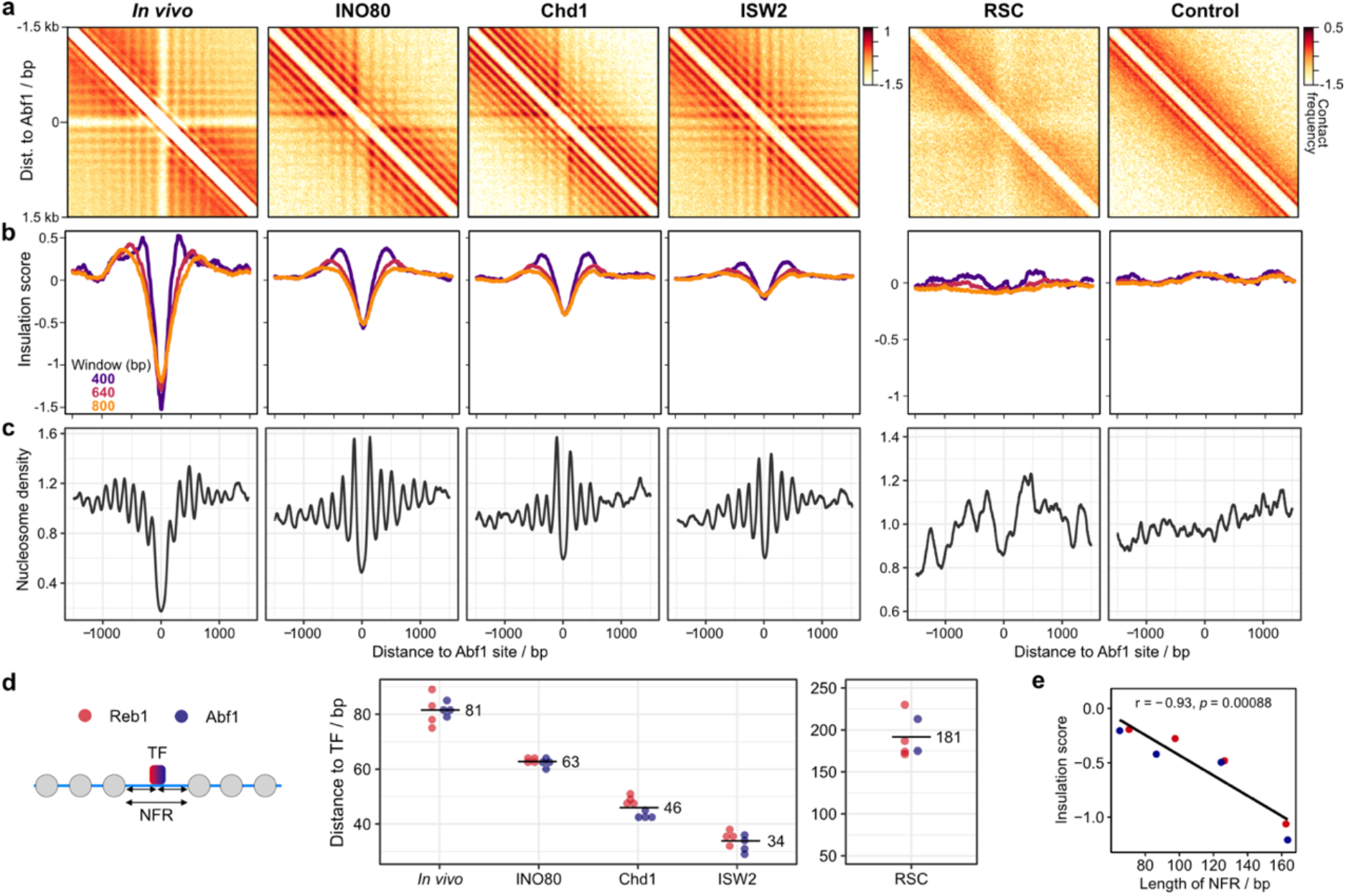
NFR width correlates with insulation strength of chromatin domains. **a,** Pile-up analyses of contact matrices aligned at Abf1 binding sites. Chromatin used for in vitro Micro-C was incubated with the indicated remodeler and the transcription factors Abf1 and Reb1 or with the transcription factors only (control). In vivo Micro-C data^21^ are shown for comparison. Log^10^ interaction counts are plotted at 20 bp resolution. **b,** Insulation scores derived from Micro-C data as described in panel a calculated at 80 bp resolution. Three different sliding windows are shown. **c,** Nucleosome occupancy profiles (MNase-seq) corresponding to panels a and b. **d,** Distances from nucleosome border to transcription factor (TF) calculated as indicated in scheme on the left. Values are derived from MNase-seq data as plotted in panel c and Extended Data Fig. 4d. Distances from two replicates and upstream or downstream direction are shown. Numbers indicate mean distances. **e,** Correlation plot of insulation score minima at Abf1 (panel b) and Reb1 centers (Extended Data Fig. 4c) versus length of nucleosome free region (NFR; panel d). Pearson’s correlation r and statistical significance p is indicated. Minima are derived from insulation scores calculated with an 800 bp window.

The meta matrices of *in vitro* chromatin reconstituted with remodelers with spacing activity closely resemble the *in vivo* matrices. However, there are subtle differences in the interaction patterns that are established by INO80, Chd1 and ISW2 with respect to insulation and boundary strength data (Fig. 3a,b, Extended Data Fig. 4b,c). INO80 generates the most distinct boundaries which create very strong insulation between the regions upstream and downstream of the Abf1 and Reb1 binding sites. Remodeling with Chd1 and ISW2 results in less insulation, with ISW2 creating the weakest domain boundaries. These differences are also apparent in the contact matrices from individual chromatin regions (Fig. 2, Extended Data Fig. 3).

To further investigate chromatin properties that regulate insulation strength, we compared the Micro-C data to MNase-seq meta profiles that show the distribution of nucleosomes in corresponding regions (Fig. 3c, Extended Data Fig. 4d). These data confirm that regular nucleosome positioning is required for the formation of chromatin domain boundaries. The MNase-seq data also show that INO80 forms the widest NFR of the three remodelers with spacing activity, followed by Chd1 and ISW2. This indicates that boundary strength is dependent on the size of the NFR established at transcription factor binding sites. To confirm this, we quantified the width of the NFR by calculating the distance between the nucleosome borders and transcription factor binding sites (Fig. 3d). We find that INO80, Chd1 and ISW2 create NFRs of distinct sizes, which vary between 63, 46, and 34 bp in width, respectively. Plotting the average boundary strength against these NFR sizes indicates that insulation strength scales proportionally with NFR width (Fig. 3e).

Consistent with the literature^24,28,30^, we find that RSC does not drive the formation of regular nucleosome arrays, but does generate very wide NFRs (Fig. 3c,e). We find that this results in a chromatin interaction pattern characterized by a depletion of signal at the NFR itself and weak insulation at the NFR. We therefore conclude that insulation strength is dependent on both the width of the NFR at the chromatin domain boundary and the formation of a regularly spaced nucleosome array, phased to the boundary site. Strong insulation *in vivo* is likely facilitated by a complex interplay between several remodelers^45,46^.

### Nucleosome linker length determines chromatin domain compaction

The *in vitro* chromatin reconstituted with the different remodelers does not only vary in NFR width, but also with regard to nucleosome linker length. INO80, ISW2 and Chd1 create linker DNA of 41 bp, 29 bp, and 22 bp, respectively (Fig. 4a). To explore how linker length relates to general compaction of chromatin domains, we plotted the interaction frequencies as a function of genomic distance for chromatin reconstituted with these three remodelers (Fig. 4b, Extended Data Fig. 5a). These interaction decay curves show a similar pattern of nucleosome interactions as previously observed in *in vivo* Micro-C and Hi-CO data^18,21,39^. The interaction decay is steepest for Chd1, which indicates that interactions between nucleosomes that are separated by large distances (>1000 bp) are relatively rare in Chd1-remodeled chromatin. Such long-range interactions are more frequent in chromatin remodeled with INO80 and ISW2. This suggest that nucleosomes with longer linker lengths form more compact chromatin. A possible explanation for these observations is that short linkers do not provide enough flexibility to support folding of chromatin into very compact structures.

**Figure 4.**
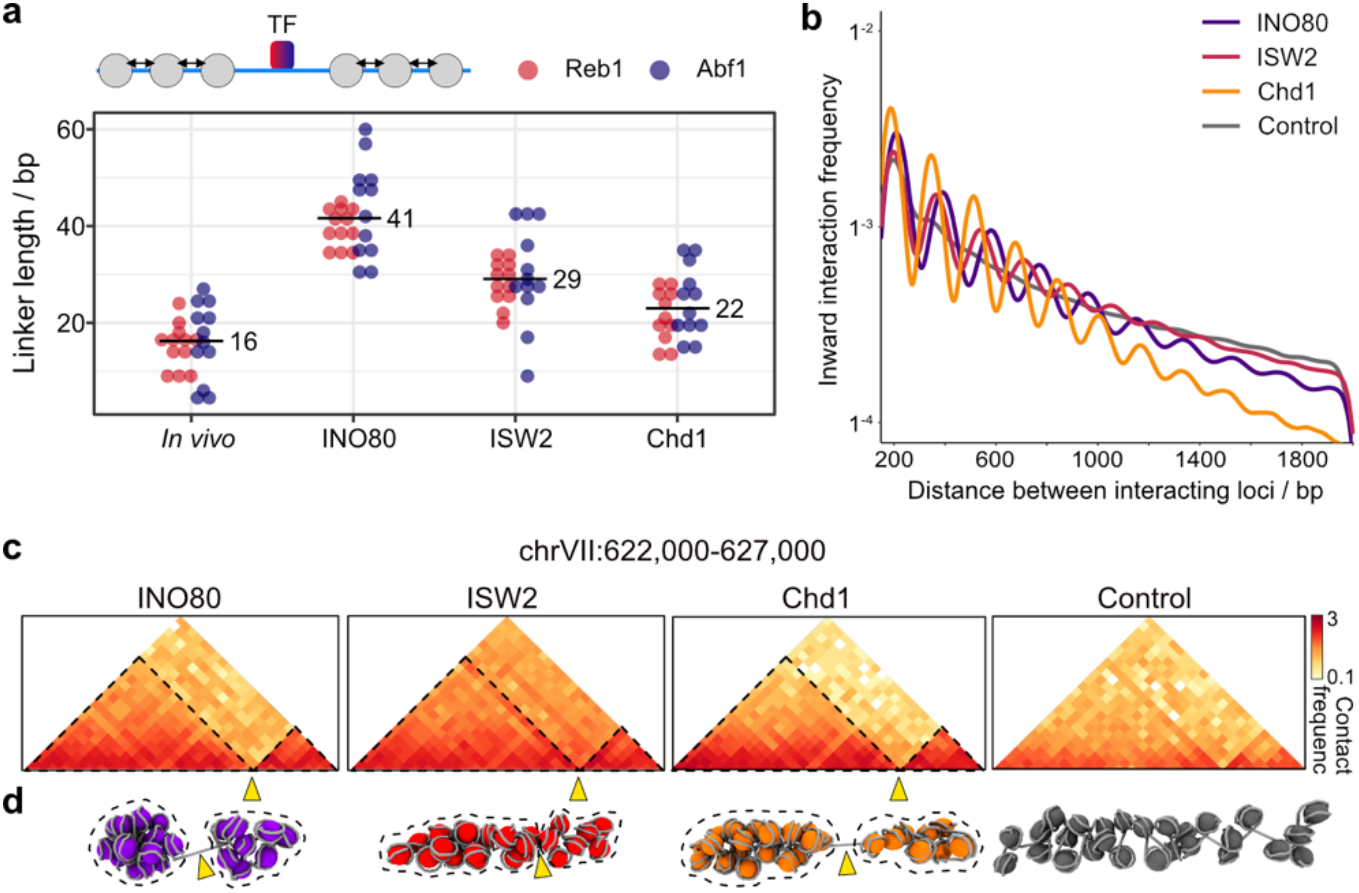
Chromatin domain compaction depends on DNA linker length. **a,** Values for linker distances plotted as indicated in the scheme on the top. Data are derived from MNase-seq as shown in Figure 3c and Extended Data Figure 4d. Three individual linker distances from upstream or downstream directions and from two replicates are plotted. Numbers indicate mean distances. **b,** Interaction frequency as a function of genomic distance plotted for inward facing nucleosome interactions derived from in vitro Micro-C data from chromatin incubated with the indicated remodeler and the transcription factors Abf1 and Reb1 or with the transcription factors only (control). **c,** Nucleosome-binned contact matrices of in vitro Micro-C data for the indicated 3 kb region. **d,** Molecular dynamics simulations of regions shown in panel c. Arrowheads points towards NFR boundaries corresponding to panel c. Stippled lines highlight chromatin domains.

To further explore higher-order folding of *in vitro* chromatin, we performed molecular dynamics simulations with a previously established pipeline^39^. These simulations are based on interaction data binned at nucleosome resolution and nucleosome orientation (Fig. 4c, Extended Data Fig. 5b). The resulting 3D models highlight the basic features of chromatin folding and their dependence on chromatin remodeling (Fig. 4d, Extended Data Fig. 5c). In the absence of remodelers, the nucleosomes do not form any specific, organized structures. However, in the presence of transcription factors and chromatin remodelers with spacing activity, the nucleosomes are organized in distinct domains that are separated by NFRs. Furthermore, comparison of the three remodelers highlights that INO80- and ISW2-remodeled chromatin with longer linkers is more compact compared to chromatin with shorter linkers that has been remodeled with Chd1. Our data are consistent with previous studies that have reported a negative relationship between nucleosome linker length and chromatin compaction^47^.

## DISCUSSION

A fundamental question in genome biology concerns the mechanisms that drive higher-order folding of chromatin and the formation of chromatin domains. Previous studies have used reconstitution of relatively small or non-natural chromatin fibers to investigate the impact of different epigenetic modulators, including histone modifications and chromatin-associated proteins, on nucleosome structure and chromatin compaction^48–50^. For the first time, we combine chromosome-wide reconstitution of native chromatin with assessment of 3D genome structure at the level of chromatin domains at very high resolution. Our unique approach has enabled us to gain fundamentally new insights into the mechanisms underlying higher-order genome folding and the interplay between genome structure and function.

By reconstituting chromatin in the presence of transcription factors and chromatin remodelers, we demonstrate that *in vivo-*like nucleosome positioning is sufficient for the establishment of native chromatin domains in *S. cerevisiae*. The boundaries of these domains are formed by wide NFRs. However, the formation of a wide NFR without regular positioning of the flanking nucleosomes, as reconstituted in the presence of the chromatin remodeler RSC, is not sufficient for the establishment of domains. We therefore conclude that the formation of chromatin domains depends on both a strong NFR and a regularly phased and spaced array of nucleosomes (Fig. 5a). In addition, we find that nucleosome linker length determines the compaction of the obtained chromatin domains, with longer linkers found in more compact chromatin (Fig. 5b).

**Figure 5.**
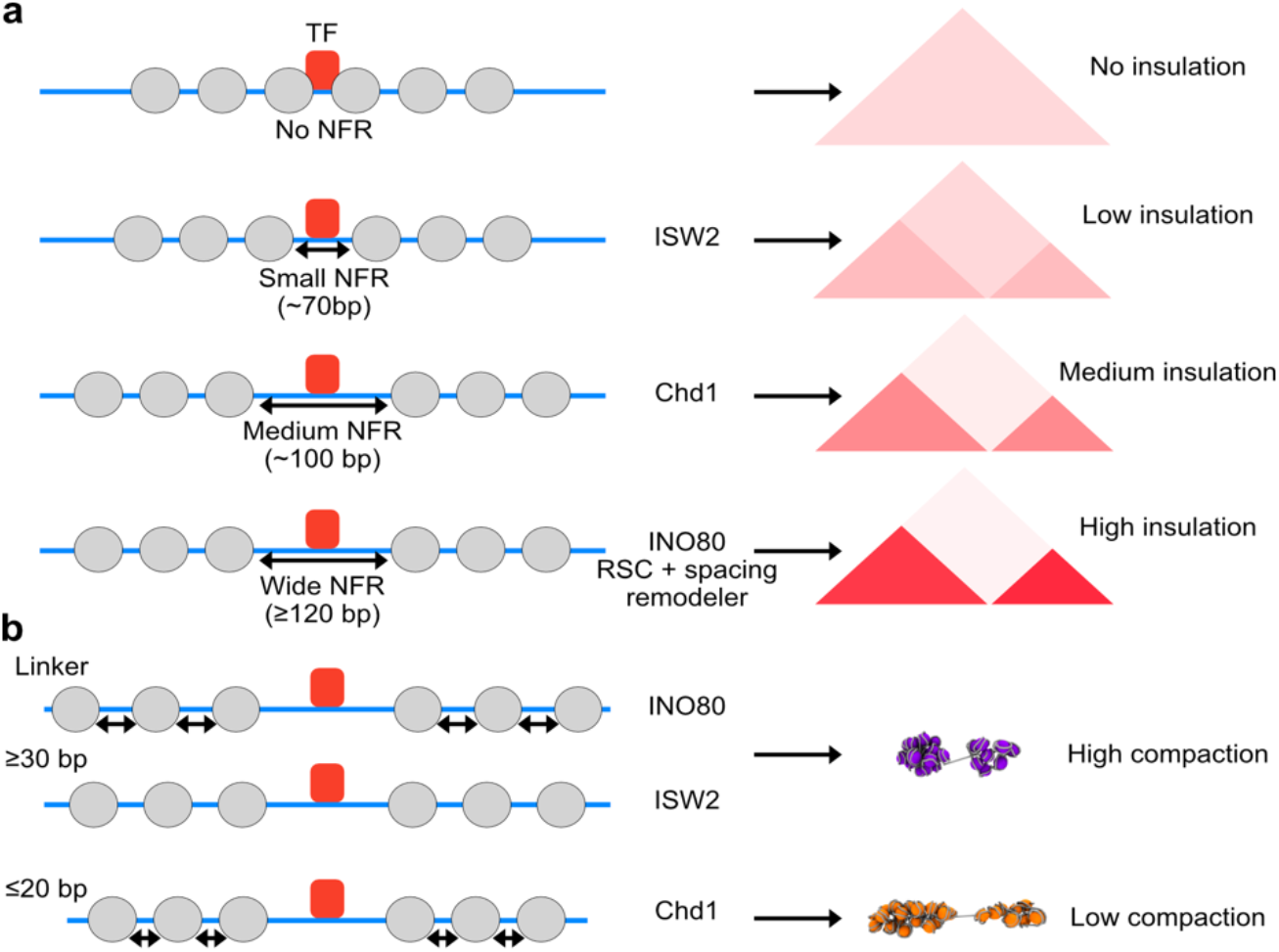
Model for the mechanism of chromatin domain formation in S. cerevisiae. **a,** Regular nucleosome positioning is sufficient for chromatin domain formation. The strength of the domain boundaries depends on the width of the nucleosome free region (NFR), which is generated in vivo by a combination of regulatory proteins, including sequence-specific transcription factors (TFs; e.g, Abf1, Reb1) and ATP-dependent chromatin remodelers (e.g., INO80, ISW2, Chd1, RSC). **b,** Compaction of chromatin domains depends on linker length, with long linkers (≥ 30 bp) leading to more compaction compared to shorter linkers. Different linker lengths are generated in vivo by the action of various ATP-dependent chromatin remodelers.

The vast majority of *in vitro* domain boundaries are formed at Abf1 and Reb1 binding sites. However, we also detect boundaries at unbound regions in which the DNA sequences are predicted to have particular shapes that can be recognized by INO80 and Chd1 (Fig. 2, Extended Data Fig. 3b). This indicates that domain formation is not dependent on the binding of a transcription factor at the boundary. We therefore conclude that physical properties of histone-DNA interactions alone are sufficient for the spontaneous formation of chromatin domains. We speculate that the separation of chromatin into two distinct domains is primarily driven by the DNA persistence length. To obtain interactions between two neighboring regions of chromatin, the DNA must bend by 180° or more. However, DNA is a very stiff polymer with a persistence length of 50 nm (~150 bp)^51^. Therefore, positively charged proteins, such as histone proteins, are required to neutralize the negative charge of DNA and enable DNA to bend^52^. This can explain why a stretch of naked DNA that is not bound by histone proteins can act as a stiff spacer and induce strong insulation between two regions of chromatin. The formation of distinct domains on either side of this boundary is likely dependent on interactions between the regularly spaced nucleosomes^53^.

Our findings have important implications for our understanding of how genome structure and function are related in eukaryotic species. For example, an important open question is whether transcription is responsible for the formation of chromatin domains or whether chromatin domains are formed first to enable transcription afterwards. In other words: “Does form follow function, or does function follow form?”^54^ Our results demonstrate that transcription is not required for the formation of chromatin domains. This may appear inconsistent with previous *in vivo* observations, since it has been reported that domain boundaries in yeast overlap with active gene promoters and that boundary strength scales with increased RNA polymerase II binding^18,21^. Although this could be interpreted as a direct role for transcription in genome organization, it is important to note that wide NFRs are a prerequisite for the formation of the pre-initiation complex (PIC) and that PIC occupancy scales with increased NFR width^55–57^. The *in vivo* observations can therefore also be explained by a pivotal role for nucleosome positioning in the regulation of both chromatin domains and transcription. This model is in line with computer simulations that have shown that nucleosome positioning alone can predict domain organization of yeast interphase chromosomes^58^. We conclude that nucleosome positioning, and not transcription, is the main driver of chromatin domain formation in yeast.

Transcription has also been implicated in the regulation of chromatin compaction. Previous *in vivo* studies in yeast have reported that active transcription is associated with higher compaction^18^. However, it has been shown that highly transcribed genes generally have short linkers^59^, which is likely mediated by the recruitment of Chd1 to actively transcribed genes^31,60–62^. It is therefore possible that the reported connection between transcription and chromatin compaction in yeast is mediated by associated changes in nucleosome spacing.

It may be that transcription is more directly involved in higher-order chromatin organization in higher eukaryotes. Indeed, many studies have suggested that active transcription plays an important role in the regulation of chromatin structure in fly^16,19,20^ and mammals^36,37,63,64^. However, our data have important implications for the current debate about the interplay between transcription and genome folding. An important line of evidence for a role of transcription in chromatin structure is based on transcription perturbation experiments. For example, experiments based on chemical inhibition of transcription or heat shock have shown that reduced transcription levels lead to changes in 3D genome organization and a rearrangement of chromatin domains in flies^19,65^. In addition, it has been shown that transcription inhibition in human cells leads to changes in fine-scale chromatin structures surrounding enhancers and promoters^36^. However, transcription elongation plays an active role in the recruitment of chromatin remodelers^60–62^ and heat shock is associated with wide-spread changes in nucleosome positioning^55^. It is therefore possible that (some of) the observed changes in these experiments could be attributed to changes in nucleosome positioning rather than reduced transcription levels. In this respect, it is of interest that it has been shown that the formation of chromatin domains during early embryonic development in flies^66^ and mice^67,68^ does not depend on transcription.

In addition to a potential direct role for transcription, it has been shown that a loop extrusion process mediated by cohesin and CTCF plays an important role in genome organization in vertebrates^69,70^. In contrast, the strong resemblance between the higher-order structure of *in vitro* reconstituted chromatin and *in vivo* yeast chromatin indicate that loop extrusion is not required for basic domain organization of the yeast genome in interphase. It should be noted though that cohesin-dependent loop extrusion plays an important role in the regulation of yeast chromatin structure during S-phase and mitosis^71–74^.

The chromatin boundaries in reconstituted chromatin predominantly correspond to Abf1 and Reb1 binding sites. These transcription factors share many features with CTCF; they function as (mild) transcriptional activators and insulators, suppress bidirectional transcription, and have pioneering activity^28–30,75,76^. It is interesting that the fine-scale interaction patterns at Abf1 and Reb1 binding sites in yeast are very similar to the patterns at CTCF binding sites in mammals^37,40^. We have previously reported that CTCF mediates local insulation in mammalian genomes in a cohesin-independent manner^40^. In light of our findings, it is likely that this is mediated by the strong influence of CTCF on nucleosome positioning^77–79^. Together, these observations indicate that the establishment of phased, regular nucleosome arrays drives the formation of (local) chromatin boundaries across eukaryotic species. Although this process alone is sufficient to drive the basic organization of the small *S. cerevisiae* genome, higher eukaryotes have evolved additional mechanisms, including loop extrusion, to structure their larger genomes. The precise interplay between nucleosome positioning, loop extrusion, and other mechanisms involved in higher-order genome folding is an exciting area to explore in more detail in future research.

## METHODS

### Genomic plasmid library

The genomic plasmid library used in this study covers ~3 Mbp of the *Saccharomyces cerevisiae* genome, corresponding to chromosomes V-IX, and contains 384 clones. This collection is part of a tiling plasmid library that was previously constructed^80^.

The library was expanded by transformation into chemically competent *dam-/dcm-* competent *Escherichia coli* cells (NEB). For every 100 μL of competent *E. coli* cells, 2 μg of DNA was added for transformation. Transformed cells were grown on large LB agar plates with kanamycin for 36 hours at 37°C, then colonies were combined into LB medium with 50 μg/ml kanamycin and grown until OD_600_ reached 2. Plasmids were extracted via a Plasmid Extraction Kit (PC2000, Macherey & Nagel).

### Expression and purification of yeast histone octamers

Yeast histone co-expression plasmids^81^ were expressed in *Escherichia coli* BL21(DE3) RIL cells. Cells were grown until OD_600_ of 0.6 in 2 x 2 L LB medium with 100 μg/mL spectinomycin, 100 μg/mL ampicillin and 20 μg/mL chloramphenicol. Expression of histones was induced by addition of 0.8 mM isopropyl 1-thio-β-d-galactopyranoside (IPTG) for 3 hours at 37°C. Cells were harvested, resuspended in 60 mL lysis buffer (20 mM Tris-Cl pH 7.6, 500 mM NaCl, 0.1 mM EDTA and 1x Protease Inhibitor Cocktail contain›ing 0.284 μg/ml leupeptin, 1.37 μg/ml pepstatin A, 0.17 mg/ml PMSF, 0.33 mg/ml benzamidine), flash frozen in liquid nitrogen and stored at −80°C.

Thawed cells were lysed by sonication on ice with a Branson Sonifier (4 x 4 minutes, 5 seconds on pulses, 5 seconds off pulses at 50% duty cycle). The lysate was cleared for 45 minutes at 89,000 g at 4°C (Sorvall LYNX 6000 centrifuge) and the supernatant was filtered with a 0.45 μM syringe filter. The cleared supernatant was loaded onto an equilibrated HiTrap Heparin 5 mL HP column (Cytiva). The column was washed with 5 column volumes (CV) of buffer A (20 mM Tris-Cl pH 7.6, 500 mM NaCl, 0.1 mM EDTA) followed by gradient elution over 14 CV of buffer A to buffer B (20 mM Tris-Cl pH 7.6, 2 M NaCl). After SDS-PAGE analysis, peak fractions were pooled and concentrated to 1 mL using a Amicon 10 kDa MWCO concentrator at 4°C. The concentrated sample was further purified by size exclusion chromatography with a Superdex 200 10/300 Increase column (Cytiva) equilibrated with 20 mM HEPES pH 7.5, 2 M NaCl. Peak fractions were pooled, concentrated and stored at −20°C in 50% glycerol.

### Expression and purification of the transcription factors Abf1 and Reb1

Abf1 or Reb1 containing expression plasmids^24^ were expressed in *Escherichia coli* BL21 (DE3) RIL cells. 2 L of Terrific Broth medium with 100 μg/mL ampicillin and 20 μg/mL chloramphenicol was inoculated with 100 mL of preculture and grown until OD 1. Cells were cooled down, protein expression was induced with 1 mM IPTG and continued at 16°C for 16 hours at 150 rpm. Cells were harvested, resuspended in Lysis Buffer (20 mM Tris-HCl pH 8 at 4°C, 500 mM NaCl, 10 mM Imidazole, 1x Protease Inhibitor (0.284 μg/ml leupeptin, 1.37 μg/ml pepstatin A, 0.17 mg/ml PMSF, 0.33 mg/ml benzamidine)), flash frozen and stored at −80°C.

For purification, cells were thawed at RT, incubated for 30 minutes on ice with 100 μg/mL Lysozyme and then sonicated for 1 minute (10 seconds on, 10 seconds off), 50% peak power. Lysate was centrifuged for 1 hour at 89,000 g, 4°C and filtered through a 0.45 μM filter. Cleared lysate was loaded onto a pre-equilibrated HisTrap HP 5 mL column (Cytiva), washed with 20 mM Tris-HCl pH 8 at 4°C, 200 mM NaCl, 20 mM Imidazole,1x Protease Inhibitor and eluted with 20 mM Tris-HCl pH 8 at 4°C, 200 mM NaCl, 250 mM Imidazole,1x Protease Inhibitor. Protein containing fractions were pooled and loaded onto a HiTrap Q HP 5 mL column, washed with 20 mM Tris-HCl pH 8 at 4°C, 200 mM NaCl, 10% Glycerol, 1 mM DTT and eluted with a gradient from 200 mM NaCl to 1 M NaCl. Protein containing fractions were pooled, concentrated using a 30 MWCO concentrator (Amicon, Merck) and subjected to gel filtration (Superdex200 10/300 Increase column (Cytiva)) in 20 mM HEPES-NaOH pH 7.5 at 4°C, 300 mM NaCl, 10% Glycerol, 1 mM DTT. Protein containing fractions were pooled, concentrated to ~30 μM, flash frozen in liquid nitrogen and stored at −80°C.

### Expression and purification of ATP-dependent chromatin remodeling enzymes

#### Chd1 and ISW2

Hi5 cells (1.2 L) were grown in ESF-921 media (Expression Systems) and infected with V1 virus for full-length Chd1 (tagged with a N-terminal 6 × His tag, followed by a MBP tag, and a tobacco etch virus protease cleavage site) or ISW2 (Spt16 tagged with an N-terminal 6 × His tag, followed by an MBP tag, and a tobacco etch virus protease cleavage site) for protein expression. Cells were grown for 72 hours at 27 °C and subsequently harvested by centrifugation (238 g, 4 °C, 30 minutes). Cell pellets were resuspended in lysis buffer (20 mM Na·HEPES pH 7.4, 300 mM NaCl, 10% (v/v) glycerol, 1 mM DTT, 30 mM imidazole pH 8.0, 0.284 μg/ml leupeptin, 1.37 μg/ml pepstatin A, 0.17 mg/ml PMSF, 0.33 mg/ml benzamidine), flash frozen and stored at −80°C.

Chd1 was purified as previously described^23^. For ISW2 purification, frozen cell pellets were thawed, lysed by sonication and cleared by centrifugation (18,000 g, 4 °C, 30 minutes and 235,000 g, 4 °C, 60 minutes). The supernatant was filtered with 0.8-μm syringe filters (Millipore) and applied onto a HisTrap HP 5 mL column (Cytiva). The column was washed first with lysis buffer, then with high salt buffer (20 mM Na·HEPES pH 7.4, 1 M NaCl, 10% (v/v) glycerol, 1 mM DTT, 30 mM imidazole pH 8.0, 0.284 μg/ml leupeptin, 1.37 μg/ml pepstatin A, 0.17 mg/ml PMSF, 0.33 mg/ml benzamidine), and finally with lysis buffer. Protein was eluted with 20 mM Na·HEPES pH 7.4, 300 mM NaCl, 10% (v/v) glycerol, 1 mM DTT, 500 mM imidazole pH 8.0 onto a prepacked Amylose column (NEB). HisTrap HP column was disconnected, the Amylose column was washed with 20 mM Na·HEPES pH 7.4, 150 mM NaCl, 10% (v/v) glycerol, 1 mM DTT and protein was eluted with wash buffer containing 4% Maltose. Protein containing fractions were pooled and subjected to TEV protease digestion for 6 h. 10 mM Imidazole was added and the sample was applied to a HisTrap HP 5 mL which was attached to a HiTrap Q HP 5 mL column. HisTrap was detached, Q column was washed and eluted with a gradient to 100% high salt buffer (20 mM Na·HEPES pH 7.4, 1 M NaCl, 10% (v/v) glycerol, 1 mM DTT). Sample was concentrated in a 50 kDa concentrator (Amicon, Merck) and loaded onto a Sepharose6 10/300 Increase pre-equilibrated in 20 mM Na·HEPES pH 7.4, 300 mM NaCl, 10% (v/v) glycerol, 1 mM DTT. Protein containing fractions were pooled, concentrated, flash frozen and stored at −80°C.

#### INO80

INO80 was purified endogenously from an overexpression strain^82^. 6 × 2 L of prewarmed YP medium with 2% Raffinose was inoculated at OD_600_ 0.1, grown for ~20 hours at 30°C at 150 rpm and induced around OD_600_ 5 with 2% Galactose (final concentration). After 3 hours induction, cells were harvested, resuspended in 100 mL of lysis buffer (50 mM HEPES-KOH pH 7.5, 500 mL KCl, 1 mM EDTA, 8 mM MgCl_2_, 0.1% NP-40, 20% Glycerol, 4 mM DTT, 0.496 μg/ml leupeptin, 2.74 μg/ml pepstatin A, 0.34 mg/ml PMSF, 0.66 mg/ml benzamidine) and frozen dropwise into liquid nitrogen. Popcorn was subjected to cryo-milling (Spex Freezer/Mill 6875D) and stored at −80°C.

All following steps were performed at 4°C unless stated otherwise. Powder was thawed in a 30°C-waterbath, then centrifuged at 30,000 g for 15 minutes and brought to a final concentration of 400 mM KCl. Lysate was subjected again to centrifugation at 90,000 g for 45 minutes. Cleared supernatant was incubated with 600 μL prewashed FLAG-M2 beads for 1 h. Beads were washed with wash buffer (25 mM HEPES-KOH pH 8, 200 mM KCl, 10% Glycerol, 0.001% IGEPAL-CA630, 2 mM MgCl_2_, 1 mM DTT) and eluted with wash buffer + 0.22 mg/mL FLAG peptides. Elution was loaded onto a HiTrap Q HP 1 mL column equilibrated with 25 mM HEPES-KOH pH 8, 200 mM KCl, 10% Glycerol, 1 mM MgCl_2_, 1 mM DTT. The column was washed and then eluted with a gradient to 1 M KCl. INO80 containing fractions were pooled, concentrated, flash frozen and stored at −80°C.

#### RSC

RSC was purified from yeast using the TAP-tag method. Rsc2-TAP-tagged yeast strain (YSC1177-YLR357W, Dharmacon) was fermented in 250 L 3% YEP broth (w/v, Formedium) supplemented with 2% glucose, 50 g L^-1^ ampicillin sodium salt and 12.5 g L^-1^ tetracycline hydrochloride until OD_600_ 10. Cells were harvested, washed with ice-cold water and resuspended 1:1 (v/v) in Lysis Buffer (50 mM K·HEPES pH 7.6 at 4°C, 700 mM KOAc, 1 mM MgCl_2_, 5% Glycerol, 1 mM DTT, 2x PI (0.56 μg/mL leupeptin, 2.74 μg/mL pepstatin A, 0.34 mg/mL phenylmethylsulfonyl fluoride, 0.66 mg/mL benzamidine)). Resuspended cells were frozen dropwise into liquid nitrogen, subjected to cryo-milling (Spex Freezer/Mill 6875D) and stored at −80°C.

All following steps were performed at 4°C unless stated otherwise. Powder was thawed in a 30°C-waterbath and slurry was first centrifuged at 14,000 g for 15 minutes followed by ultracentrifugation at 200,000 g for 105 minutes. Clear phase was taken and supplemented with 50 mM KOAc. Then, the lysate was incubated with prewashed IgG Sepharose 6 Fast Flow resin (Cytiva) for 3 h. Resin was collected by centrifugation and washed 2x with Wash buffer (40 mM K·HEPES pH 7.6, 250 mM KAc, 10% Glycerol, 0.5 mM DTT, 10 mM EDTA pH 8, 1x PI) and 1x with TEV-Elution Buffer (40 mM K·HEPES pH 7.6, 200 mM KAc, 10% Glycerol, 0.5 mM DTT, 10 mM EDTA pH 8). Protein was eluted by addition of TEV-Elution buffer containing 10 μg/mL TEV protease and incubation for 2 hours at 15°C with gentle agitation. Elution was loaded onto a pre-equilibrated 1 mL HiTrap Q HP column (Cytiva), washed and eluted over 40 CV from Buffer A to B (Buffer A: 40 mM HEPES-KOH pH 7.6, 200 mM KAc, 10% Glycerol, 1 mM DTT, Buffer B: 40 mM K·HEPES, 1.5 M KAc, 10% Glycerol, 1 mM DTT). RSC-containing fractions were pooled, concentrated, flash frozen and stored at −80°C.

### *In vitro* reconstitution of yeast chromatin

#### Salt gradient dialysis (SGD)

All steps were performed at room temperature. 46 μg of plasmid library DNA were mixed with 40 μg of recombinant *S. cerevisiae* histone octamers in 400 μL assembly buffer (10 mM HEPES-NaOH pH 7.6, 2 M NaCl, 0.5 mM EDTA pH 8, 0.05% IGEPAL CA-630, 0.2 μg/μL BSA). Sample was transferred to a Slide-A-Lyzer MINI dialysis cup (3.5 MWCO, ThermoFisher Scientific), which was placed in a 3 L beaker containing 300 mL high salt buffer (10 mM HEPES-NaOH pH 7.6, 2 M NaCl, 1 mM EDTA, 0.05% IGEPAL CA-630). Sample was gradually dialyzed against a total of 3 L low salt buffer (10 mM HEPES-NaOH pH 7.6, 50 mM NaCl, 0.5 mM EDTA, 0.05% IGEPAL CA-630) while stirring via a peristaltic pump over 12-14 h. After complete transfer of low salt buffer, samples were dialyzed against 1 L low salt buffer for 1 h. The chromatin was centrifuged for 1 minute at 10,000 rpm and the supernatant was transferred to a fresh protein low-binding tube. We refer to the *in vitro*-reconstituted chromatin as “SGD chromatin”. DNA concentration of the SGD chromatin was measured using a spectrophotometer (Nanodrop) and it was stored for maximum 4 weeks at 4°C.

#### Remodeling of SGD chromatin

Remodeling reactions contained 1) 30 μL of SGD chromatin 2) transcription factors Abf1 and Reb1 and 3) one of the chromatin remodeling enzymes, in remodeling buffer (30 mM HEPES-KOH pH 7.6, 3 mM MgCl_2_, 2.5 mM ATP, 1.25 mM TCEP, 0.4 mM DTT, 10 mM creatine phosphate, 0.25 mM EGTA, 0.15 mM EDTA, 80 mM KOAc, 15 mM NaCl, 0.015% IGEPAL CA-630, 16.5% glycerol, 60 μg/mL BSA, 10 μg/mL creatine kinase) for a total reaction volume of 100 μL. The SGD chromatin was generated with the transcription factors, Abf1 and Reb1, alone (final concentration of 50 nM each) or with the transcription factors and one of the following remodeling enzymes: INO80, Chd1, ISW2 or RSC at a final concentration of 24 nM or 12 nM for RSC. Chd1-remodeling was performed in in remodeling buffer containing only 50 mM KOAc. Different replicates represent different batches of reconstituted chromatin.

### *In vitro* Micro-C

All crosslinking steps were performed at 30°C. First, remodeled SGD chromatin was incubated with disuccinimidyl glutarate (DSG) at a final concentration of 0.75 mM for 20 minutes. Then, formaldehyde (FA) was added at a final concentration of 0.05% for another 10 minutes. The crosslinking reactions were quenched with 10x Quenching Buffer (100 mM Tris pH 7.5, 80 mM aspartate, 20 mM lysine) for 15 minutes. Crosslinked chromatin was diluted in digestion buffer (10 mM Tris-Cl pH 7.6, 1.7 mM CaCl2) and supplemented with 0.4 Ku/μL (final concentration) of MNase (New England Biolabs). The reaction was terminated with 2x STOP-Buffer (10 mM Tris-HCl pH 7.5, 20 mM EDTA, 400 mM NaCl, 2 mM EGTA) for 5 minutes.

The crosslinked and MNase-digested chromatin was purified using an Amicon Ultra 50 kDa centrifugal filter to remove <100 bp DNA fragments. The chromatin was diluted with low salt buffer (20 mM HEPES-NaOH, 50 mM NaCl, 0.5 mM EDTA, 0.05% IGEPAL CA-630), loaded into the filter and the reaction was centrifuged (2000 g, 2 minutes, room temperature). The reaction was washed with low salt buffer two more times and concentrated to ~100 μL.

The proximity ligation reaction was adapted from the previously described MCC protocol^41^, in which DNA end repair, phosphorylation and ligation are performed in a single tube. Nucleosomes were resuspended in T4 Ligation buffer supplemented with 0.4 mM dNTP, 2.5 mM EGTA, 20 U/ml T4 Polynucleotide Kinase, 10 U/ml DNA Polymerase I Large (Klenow) Fragment and 30 U/ml T4 DNA ligase in a total reaction volume of 400 μL. The reaction was incubated at 37°C for 2 hours followed by 22°C for 8 hours at 300 rpm on an Eppendorf Thermomixer. The chromatin was reverse crosslinked with the addition of 1 mg/mL Proteinase K (Life Technologies) and 0.5% SDS at 65°C for >16 hours. For DNA extraction, the reaction was supplemented with glycogen and subjected to ethanol precipitation. Digestion and ligation efficiencies were assessed using Fragment Analyzer. A successful ligation is indicated by the increase in fragment sizes of > 320 bp (Extended Data Fig. 2e). Unligated mono-nucleosome fragments were further removed by purification with AMPure XP beads in a 0.9:1 ratio.

Approximately 150-200 ng of each library was sonicated to a mean fragment size of 200 bp using a Covaris S220 Ultrasonicator (peak incident power 175; duty factor 10%; cycles per burst 200; treatment time 250 s) followed by purification with AMPure XP beads in a 1.8:1 ratio. 170-200 ng of library was indexed using NEBNext Ultra II DNA Library Prep Kit for Illumina (New England Biolabs). The manufacturer’s protocol was followed with the following deviations: to maximize library complexity and yield, the PCR was performed in duplicate per ligation reaction using Herculase II reagents (Agilent Technologies). The parallel library preparations and PCR reactions were subsequently pooled for each reaction. The quality of the library and the molar concentration were measured using Fragment Analyzer and Qubit. The material was sequenced on Illumina NextSeq 550 with 150 bp paired end reads, with each replicate having ~30 million reads.

### MNase-seq

10 μL crosslinked, MNase-digested and purified chromatin was used to assess the quality of MNase digestion. After digestion, DNA was extracted by ethanol precipitation. Fragment sizes were evaluated on a Fragment Analyzer. Approximately 15-20 ng of the digestion control was indexed using NEB Next Ultra II Library Kit using the manufacturer’s protocol. The material was sequenced on Illumina NextSeq 550 with 40 bp or 150 bp paired end reads.

### Reference Datasets

*In vivo* Micro-C data^21^ were downloaded from GEO (GSM2262329, GSM2262330, GSM2262331) and re-analyzed using HiC-Pro^83^. Briefly, reads were mapped to the SacCer3 reference genome using Bowtie2^84^, and valid pairs were extracted. The list of valid pairs for replicates was merged.

ChIP-exo data^28^ (Abf1: GSM4449154; Reb1: GSM4449823) were mapped against the SacCer3 genome using Bowtie^85^. 5’ ends of reads were used and extended to 3 bp. Coverage files were calculated in R using GenomicAlignments^86^ and visualized in the integrated genome viewer^87^ (igv 2.8.6) for single loci analysis. For alignments, the corresponding called ChIP-exo peaks from http://www.yeastepigenome.org/ were intersected with corresponding PWM motifs that were previously generated^23^. This resulted in 119 sites for Abf1 and 128 sites for Reb1 on chromosomes V-IX.

### Analysis of *in vitro* Micro-C data

#### Mapping

Analysis was performed using the MCC pipeline as previously described^41^. Briefly, adapter sequences were removed using Trim Galore (Babraham Institute, v.0.3.1) and paired-end reads were reconstructed using FLASH^88^ (v.1.2.11). Reconstructed reads were mapped to the ~3 Mbp plasmid library with the non-stringent aligner BLAT^89^. Uninformative reads (e.g. plasmid backbone) were discarded, while the mapped reads in the FASTQ files were further mapped to the SacCer3 reference genome using Bowtie2^84^. The aligned reads were then processed to identify the ligation junction and remove PCR duplicates using the MCC pipeline available from https://process.innovation.ox.ac.uk/software/p/16529a/micro-capture-c-academic/1). The MCC pipeline generates a list of the chimeric pairs, the base pair coordinates of the ligation junction, and direction of the read (upstream or downstream of the ligation junction). The numbers of reads at each process are summarized in Supplementary Tables 1 and 2. Interactions that are inter-chromosomal and less than 147 bp apart were removed from the pairwise interaction list. The read pair orientation was classified into four groups based on the direction of the read relative to the junction (inward, outward, tandem-entry, tandem-exit) as described previously^39^. The ligation junction was further shifted by 80 bp to the nucleosome dyad.

#### Contact matrices

To generate contact matrices, chimeric pairs were aggregated in the cooler format using the cooler package^90^ at 20, 40 and 80 bp resolution. Pearson correlation of 80 bp resolution contact matrices of replicate 1 and 2 was calculated using the hicCorrelate function of hiCExplorer package^91^. The chimeric pairs list for the two replicates were combined and these data are represented throughout the manuscript. To create the nucleosome-binned contact matrix, we assigned each of the interacting pairs to one of the corresponding 66,360 nucleosome loci previously determined through MNase-seq^92^.

#### Pile-up analysis of contact matrices

Contact frequencies with 20 bp resolution were extracted within a 3000 bp window around Abf1 and Reb1 binding sites^28^ and then aggregated and visualized using the cooltools package^93^.

#### Insulation scores and boundary calling

Diamond insulation scores for 80 bp resolution contact matrices were calculated with different window size (400, 640 and 800 bp) using cooltools^93^. Insulating loci were called using a local minima detection procedure based on peak prominence. Highly insulating regions that correspond to strong boundaries are called according to the thresholding method “Li” from the image analysis field^94^. Insulation scores were averaged in a 3000 bp window around Abf1 and Reb1 binding sites^28^.

#### Interaction decay curves

Interaction counts were log^10^ transformed and plotted as a function of genomic distance for each read-pair orientation.

### Molecular dynamics simulation

3D chromatin models derived from the *in vitro* Micro-C data were generated from simulated annealing (SA)-molecular dynamics (MD) simulation as previously described^39,95^. In brief, the simulation represents nucleosomes as composed of histone and DNA beads, equivalent in number to the naturally occurring nucleosome loci. A histone octamer corresponds to four histone beads having a 3 nm radius^96^. One bead of DNA corresponds to 5.88 bp and a total of 23 sequential DNA beads is wrapped around four histone particles in a left-handed superhelical geometry for 1.65 turns with a radius of 4.18 nm and a pitch of 2.39 nm^96^.

### Analysis of MNase-seq data

For samples sequenced with 150-bp paired ends, sequencing reads were trimmed to 75 bp. For 40-bp paired-end reads, trimming was omitted. Reads were mapped with Bowtie^85^ to the *S. cerevisiae* genome SacCer3 omitting multiple matches. Mapped data were imported into R Studio using GenomicAlignments^86^ and only fragments with 125-205 bp length were kept. Nucleosome dyad length was reduced to 50 bp and smoothed by a 20 bp rolling window. Genome coverage was calculated. For single-gene visualization, data were converted to a GenomicRanges format, exported as bigwig file and loaded into the integrated genome viewer^87^ (igv 2.8.6).

Views of coverage files were generated with a 2001 bp or 3001 bp window around *in vivo* + 1 nucleosome positions^23^ or Abf1 and Reb1 binding sites^28^, respectively. Only sites within chromosomes V-XI were used (1184 sites for *in vivo* +1 nucleosome, 119 sites for Abf1, 128 sites for Reb1). Nucleosome signal was normalized per window.

Windows around *in vivo* + 1 nucleosome sites were sorted by Abf1 and Reb1 signal in the NFR. To this end, Abf1 (GSM2916412) and Reb1 (GSM2916410) ChIP-seq data^97^ were merged, aligned as described above to *in vivo* +1 nucleosomes and sorted by decreasing signal strength in a 180 bp window 160 bp upstream of the *in vivo* +1 nucleosome site.

For composite plots, mean of normalized nucleosome signal was calculated and plotted for transcription factor bound genes (top 20% of genes), transcription factor unbound genes (bottom 80 % of genes) or Abf1 and Reb1 binding sites.

Linker and NFR values were calculated as previously described^23^ In brief, peaks of nucleosome occupancy profiles around Abf1 and Reb1 binding sites were called. Then, the nucleosome repeat length was calculated (distance from peak to next neighboring peak) and 147 bp was subtracted to obtain linker length of first three nucleosomes upstream and downstream of the transcription factor binding site. For the calculation of the distance to the transcription factor binding site, the peak of the first nucleosome was called, the distance to the alignment point was calculated and 73 bp were subtracted.

## Data availability

Raw sequencing data and processed data are available for download at http://www.ncbi.nlm.nih.gov/geo/ via GEO accession number GSE220647.

## ACKNOWLEDGEMENTS

We thank Lucas Farnung for sharing purified Chd1, plasmids and advice regarding protein purification; Philipp Korber for sharing the yeast plasmid library and Abf1/Reb1 expression plasmids; Christoph Kurat and John Diffley for sharing the INO80-FLAG strain; Martin Singleton for sharing *S. cerevisiae* histone co-expression plasmids; Aleksandra Galitsyna for support with analysis of Hi-CO data; and Masae Ohno and Yuichi Taniguchi for support with molecular dynamics simulations. We are grateful to Christian Dienemann, Michael Lidschreiber, Johannes Söding, and James Walshe for feedback on the manuscript. We also thank all members of the Oudelaar and Cramer laboratories for helpful discussions. P.C. is supported by the Max Planck Society, the Deutsche Forschungsgemeinschaft (EXC 2067/1 - 390729940), and the European Research Council (Advanced Grant CHROMATRANS; 882357). A.M.O. is supported by the Max Planck Society and the Deutsche Forschungsgemeinschaft.

