## Supplementary Information for "*In vitro* reconstitution of chromatin domains"

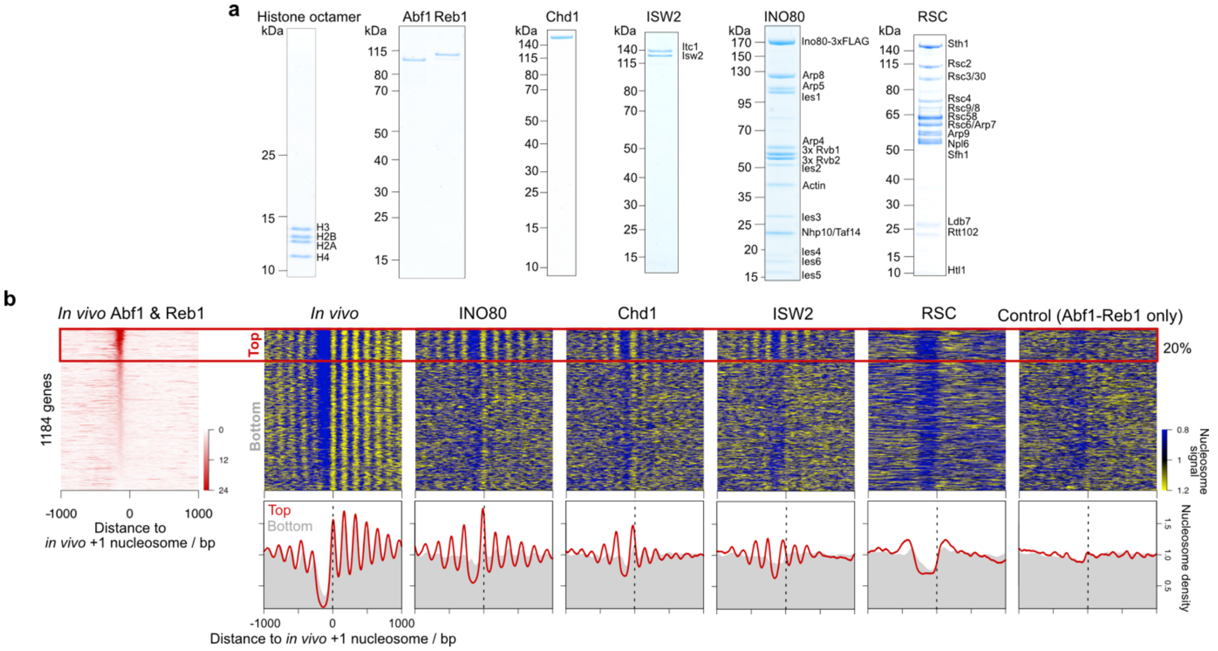


**Extended Data Figure 1. Reconstitution of native nucleosome positioning with purified proteins. a,** SDS-PAGE analysis of purified proteins used for *in vitro* chromatin reconstitution. **b,** Heat map of merged Abf1 and Reb1 SLIM-ChIP data[97] aligned at *in vivo* +1 nucleosome positions and sorted by transcription factor binding signal in nucleosome free regions (NFRs) (left). Heat maps (top) and composite plots (bottom) of nucleosome occupancy of *in vitro*-reconstituted chromatin incubated with Abf1 and Reb1 alone (control) or additionally with the indicated remodeler. Genes in heat map are sorted according to transcription factor binding as shown on the left. *In vivo* nucleosome positioning is derived from Micro-C data[21]. Composite plots show averaged nucleosome occupancy of transcription factor bound genes (red, corresponding to top 20% of genes) or transcription factor unbound genes (grey, corresponding to bottom of heat map).

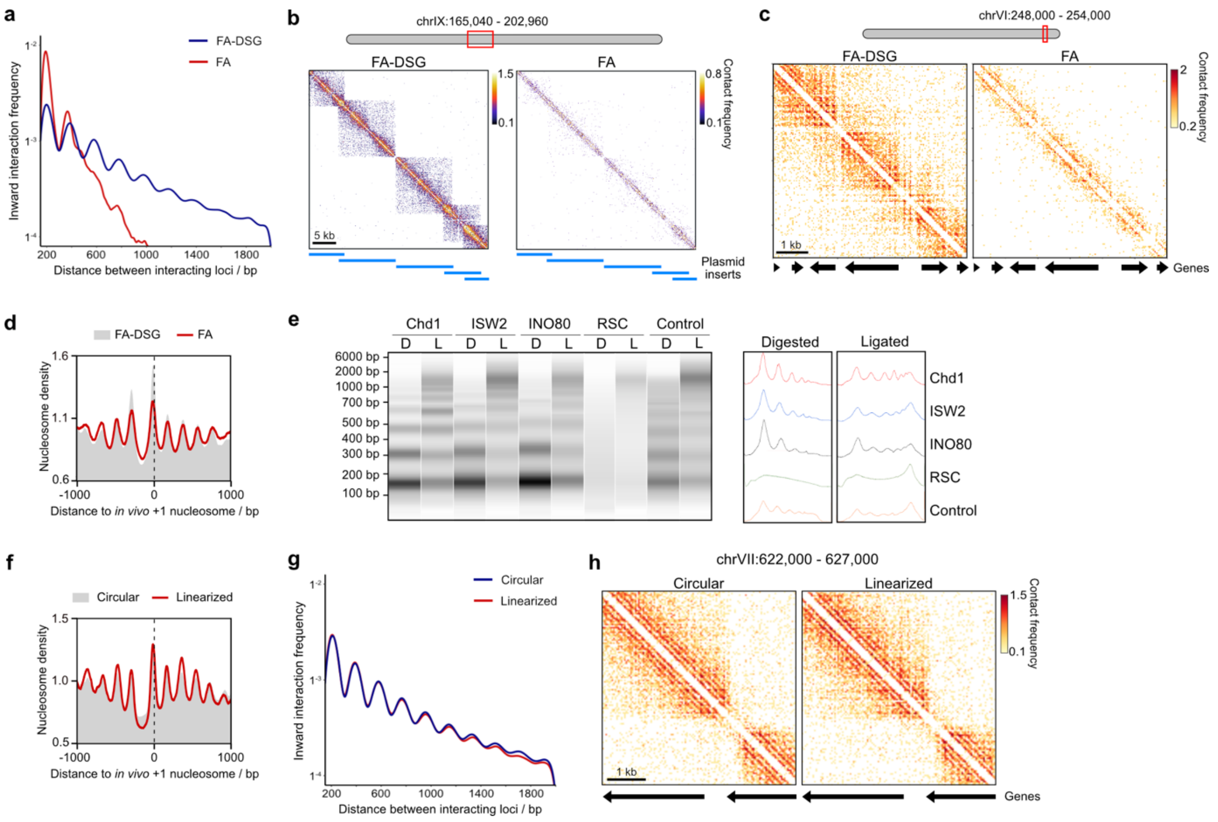


**Extended Data Figure 2. Establishment of the *in vitro* Micro-C procedure. a,** Distance decay curve of inward-facing nucleosome interactions for indicated crosslinking conditions of *in vitro* Micro-C of SGD chromatin generated in the presence of INO80, Abf1 and Reb1. 0.05% formaldehyde (FA) and 0.75 mM disuccinimidyl glutarate (DSG) was used. **b,** Contact matrices of a 37.92 kb region of *in vitro* Micro-C from chromatin samples described in panel a. **c,** Contact matrices of a 6 kb region of *in vitro* Micro-C from chromatin samples as described in panel a. **d,** Nucleosome occupancy profiles of *in vitro*-reconstituted chromatin from chromatin samples as described in panel a. **e,** Capillary gel electrophoresis of *in vitro* Micro-C samples after MNase-digestion (D) and after proximity ligation (L). *In vitro* chromatin was incubated with Abf1 and Reb1 alone (control) or with the indicated chromatin remodeler. **f,** Nucleosome occupancy profiles of *in vitro*-reconstituted chromatin incubated with INO80, Abf1 and Reb1 alone (circular) or additionally with the restriction enzyme BamHI (linearized). **g,** Distance decay curve of inward-facing nucleosome interactions of *in vitro* Micro-C from chromatin samples as described in panel g. **h,** Contact matrices of a 5 kb region of *in vitro* Micro-C data as described in panel g.

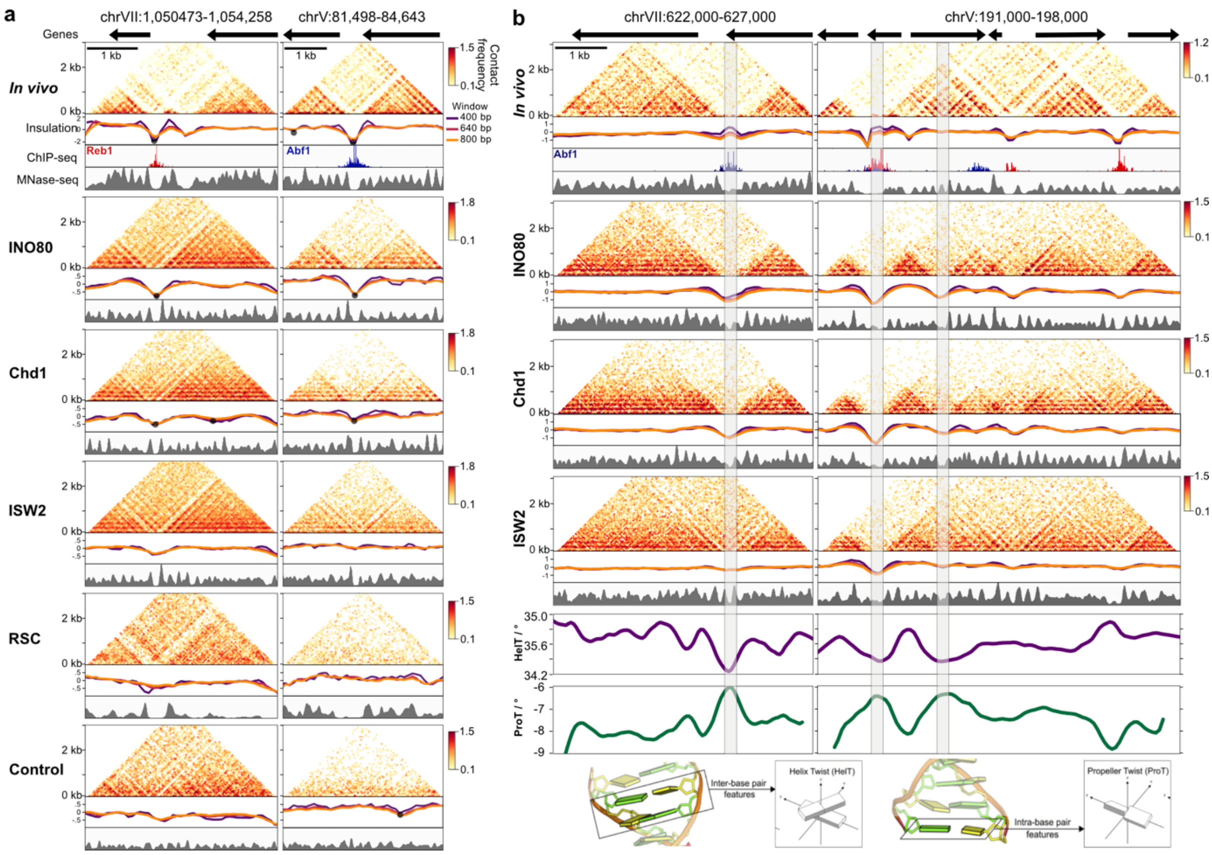


**Extended Data Figure 3. A subset of *in vitro* reconstituted domain boundaries overlaps with DNA-shape features. a,** Contact matrices, insulation scores, nucleosome occupancy profiles (MNase-seq), and Abf1 and Reb1 ChIP-exo data for the indicated regions and conditions, as described in Figure 2. **b**, DNA shape feature calculation of Helix Twist (HelT) and Propellor Twist (ProT)[44] of two regions shown in Figure 2. Grey bars highlight regions with high HelT and low ProT which overlap with strong boundaries observed *in vitro*.

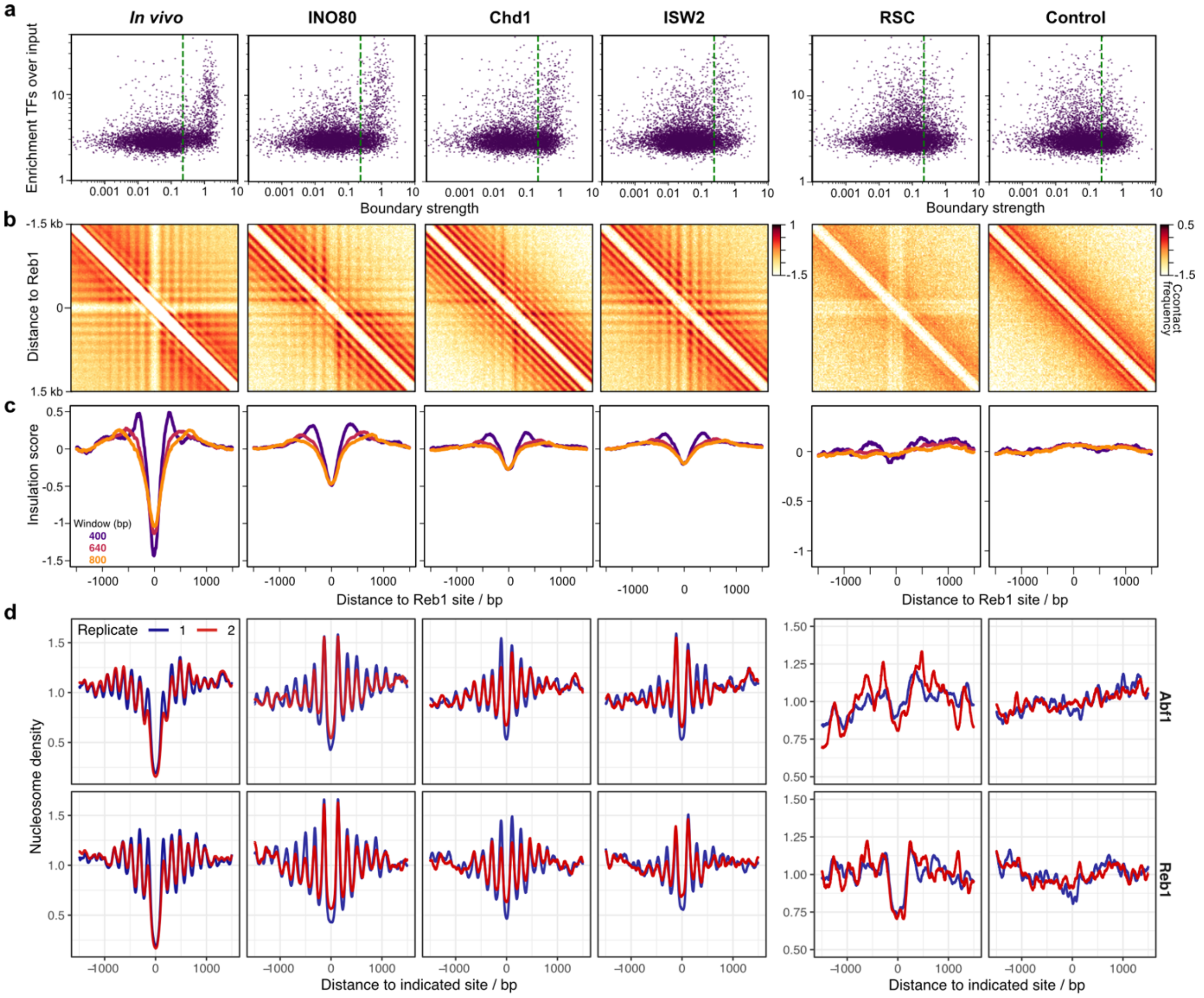


**Extended Data Figure 4. Strong domain boundaries are enriched at transcription factor binding sites which correlate with regular nucleosome positioning. a,** Transcription factor enrichment (derived from Abf1 and Reb1 ChIP-exo data[28]) as a function of boundary strength defined by insulation scores plotted with 800 bp sliding windows derived from *in vivo*[21] and *in vitro* Micro-C data. Chromatin used for *in vitro* Micro-C was incubated with the indicated remodeler and the transcription factors Abf1 and Reb1 or with the transcription factors only (control). Green dashed lines denote strong boundaries based on Li automated thresholding criteria[94]. **b,** Pile-up analysis of contact matrices aligned at Reb1 binding sites. Chromatin used for *in vitro* Micro-C was incubated with the indicated remodeler and the transcription factors Abf1 and Reb1 or with the transcription factors only (control). *In vivo* Micro-C data[21] are shown for comparison. $Log^{10}$ interaction counts are plotted at 20 bp resolution. **c,** Insulation scores of Micro-C data as described in panel b calculated at 80 bp resolution. Three different sliding windows are shown. **d,** Nucleosome occupancy profiles (MNase-seq) of individual replicates aligned at Abf1 or Reb1 binding sites corresponding to panels b and c.

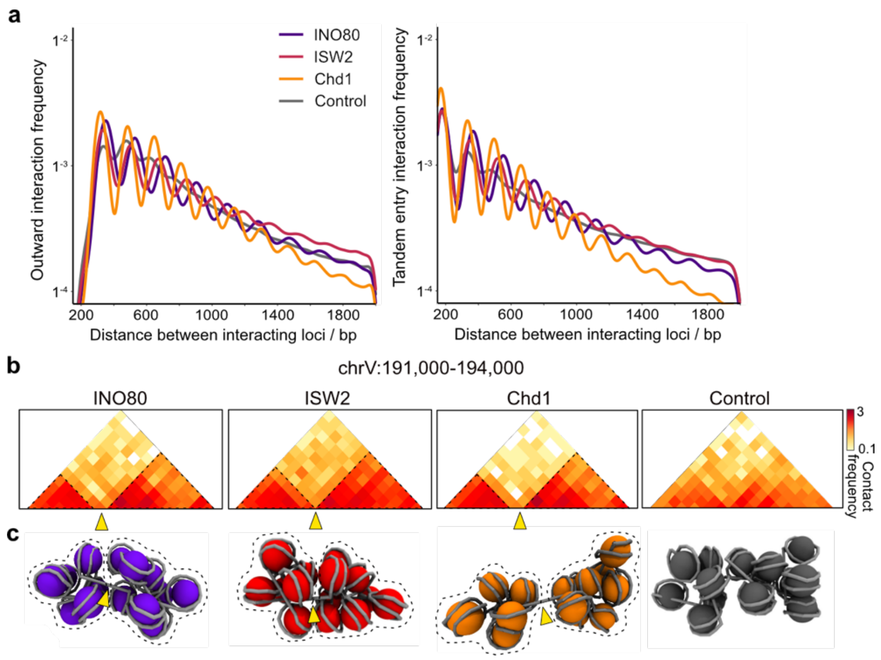


**Extended Data Figure 5. Chromatin domain compaction increases with longer linker lengths. a,** Interaction frequency as a function of genomic distance plotted for outward and tandem interactions of nucleosomes as derived from *in vitro* Micro-C data from chromatin incubated with the indicated remodeler and the transcription factors Abf1 and Reb1 or with the transcription factors only (control). **b,** Nucleosome-binned contact matrices of *in vitro* Micro-C data for the indicated region. **c,** Molecular dynamics simulations of regions shown in panel b. Arrowheads points towards NFR boundaries corresponding to panel b. Stippled lines highlight chromatin domains.
